## Supplementary Figures and Tables for "*fmo-4* promotes longevity and stress resistance via ER to mitochondria calcium regulation in *C. elegans*"

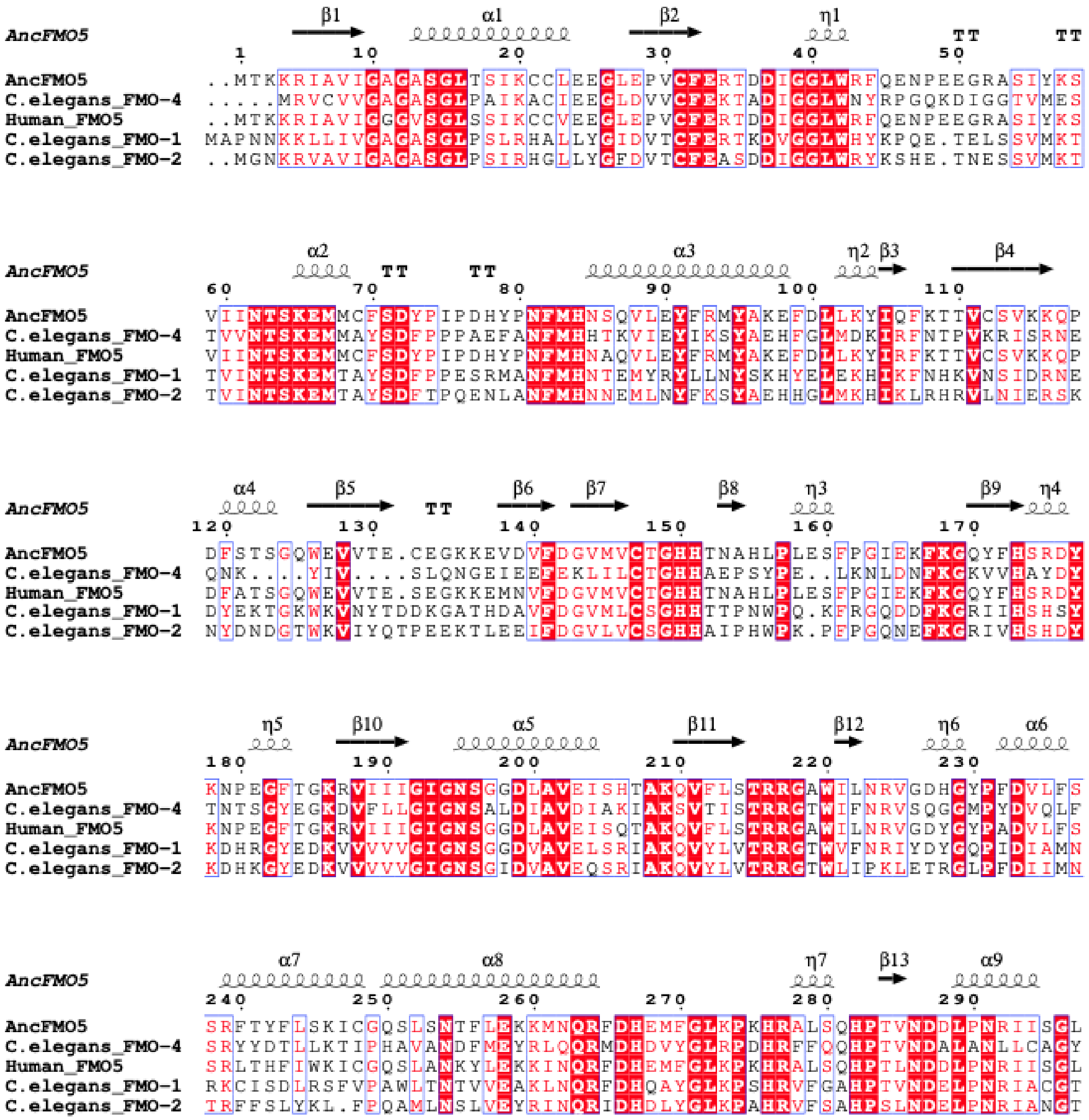
AncFMO5
β1 α1 β2 η1 TT TT
1 10 20 30 40 50
AncFMO5 ..MTKKRIAVIGAGASGLTSIKCCLEEGLEPVCFERTDDIGGLWRFQENPEEGRASIYKS
C.elegans_FMO-4 .....MRVCVVGAGASGLPAIKACIEEGLDVVCFEKTADIGGLWNYRPGQKDIGGTVMES
Human_FMO5 ..MTKKRIAVIGGGVSGLSSIKCCVEEGLEPVCFERTDDIGGLWRFQENPEEGRASIYKS
C.elegans_FMO-1 MAPNNKKLLIVGAGASGLPSLRHALLYGIDVTCFERTKDVGGLWHYKPQE.TELSSVMKT
C.elegans_FMO-2 ..MGNKRVAVIGAGASGLPSIRHGLLYGFDVTCFEASDDIGGLWRYKSHE.TNESSVMKT
AncFMO5
α2 TT TT α3 η2 β3 β4
60 70 80 90 100 110
AncFMO5 VIINTSKEMMCFSDYPIPDHYPNFMHNSQVLEYFRMYAKEFDLLKYIQFKTTVCSVKKQP
C.elegans_FMO-4 TVVNTSKEMMAYSDFPPPAEFANFMHHTKVIEYIKSYAEHFGLMDKIRFNTPVKRISRNE
Human_FMO5 VIINTSKEMMCFSDYPIPDHYPNFMHNAQVLEYFRMYAKEFDLLKYIRFKTTVCSVKKQP
C.elegans_FMO-1 TVINTSKEMTAYSDFPPESRMANFMHNTEMYRYLLNYSKHYELEKHIKFNHKVNSIDRNE
C.elegans_FMO-2 TVINTSKEMTAYSDFTPQENLANFMHNNEMLNYFKSYAEHHGLMKHIKLRHRVLNIERSK
AncFMO5
α4 β5 TT β6 β7 β8 η3 β9 η4
120 130 140 150 160 170
AncFMO5 DFSTSGQWEVVTE.CEGKKEVDVFDGVMVCTGHHTNAHLPLESFPGIEKFKGQYFHSRDY
C.elegans_FMO-4 QNK....YIV....SLQNGEIEEFEKLILCTGHHAEPSYPE..LKNLDNFKGKVVHAYDY
Human_FMO5 DFATSGQWEVVTE.SEGKKEMNVFDGVMVCTGHHTNAHLPLESFPGIEKFKGQYFHSRDY
C.elegans_FMO-1 DYEKTGKWKVNYTDDKGATHDAVFDGVMLCSGHHTTPNWPQ.KFRGQDDFKGRIIHSHSY
C.elegans_FMO-2 NYDNDGTWKVIYQTPEEKTLEEIFDGVLVCSGHHAIPHWPK.PFPGQNEFKGRIVHSHDY
AncFMO5
η5 β10 α5 β11 β12 η6 α6
180 190 200 210 220 230
AncFMO5 KNPEGFTGKRVIIIGIGNSGGDLAVEISHTAKQVFLSTRRGAWILNRVGDHGYPFDVLFS
C.elegans_FMO-4 TNTSGYEGKDVFLLGIGNSALDIAVDIAKIAKSVTISTRRGTWIFNRVSQGGMPYDVQLF
Human_FMO5 KNPEGFTGKRVIIIGIGNSGGDLAVEISQTAKQVFLSTRRGAWILNRVGDYGYPADVLFS
C.elegans_FMO-1 KDHRGYEDKVVVVVGIGNSGGDVAVELSRIAKQVYLVTRRGTWVFNRIYDYGQPIDIAMN
C.elegans_FMO-2 KDHKGYEDKVVVVVGIGNSGIDVAVEQSRIAKQVYLVTRRGTWLIPKLETRGLPFDIIMN
AncFMO5
α7 α8 η7 β13 α9
240 250 260 270 280 290
AncFMO5 SRFTYFLSKICGQSLSNTFLEKKMNQRFDHEMFGLKPKHRALSQHPTVNDDLPNRIISGL
C.elegans_FMO-4 SRYYDTLLKTIPHAVANDFMEYRLQQRMDHDVYGLRPDHRFFQQHPTVNDALANLLCAGY
Human_FMO5 SRLTHFIWKICGQSLANKYLEKKINQRFDHEMFGLKPKHRALSQHPTLNDDLPNRIISGL
C.elegans_FMO-1 RKCISDLRSFVPAWLTNTVVEAKLNQRFDHQAYGLKPSHRVFGAHPTVNDELPNRIACGT
C.elegans_FMO-2 TRFFSLYKL.FPQAMLNSLVEYRINQRIDHDLYGLKPAHRVFSAHPSLNDELPNRIANGT

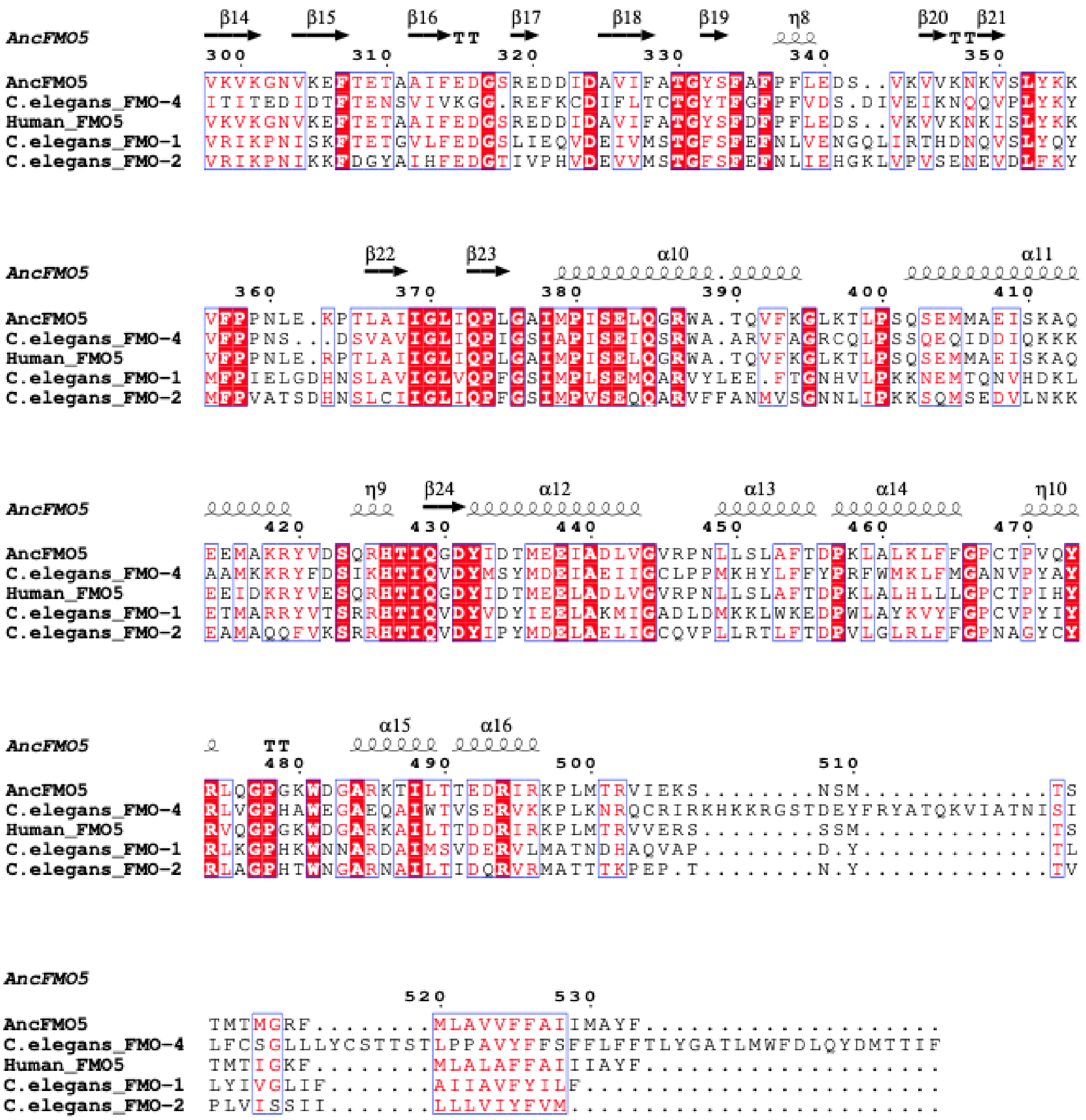


**Supplementary Figure 1. Full-length alignment of AncFMO5, Human FMO5, and *C. elegans* FMO-1, FMO-2, and FMO-4.** Alignment was conducted using Clustal Omega and ESPript 3.0 to determine an 88% overlap in catalytic residues between *C. elegans* FMO-2 and FMO-4.

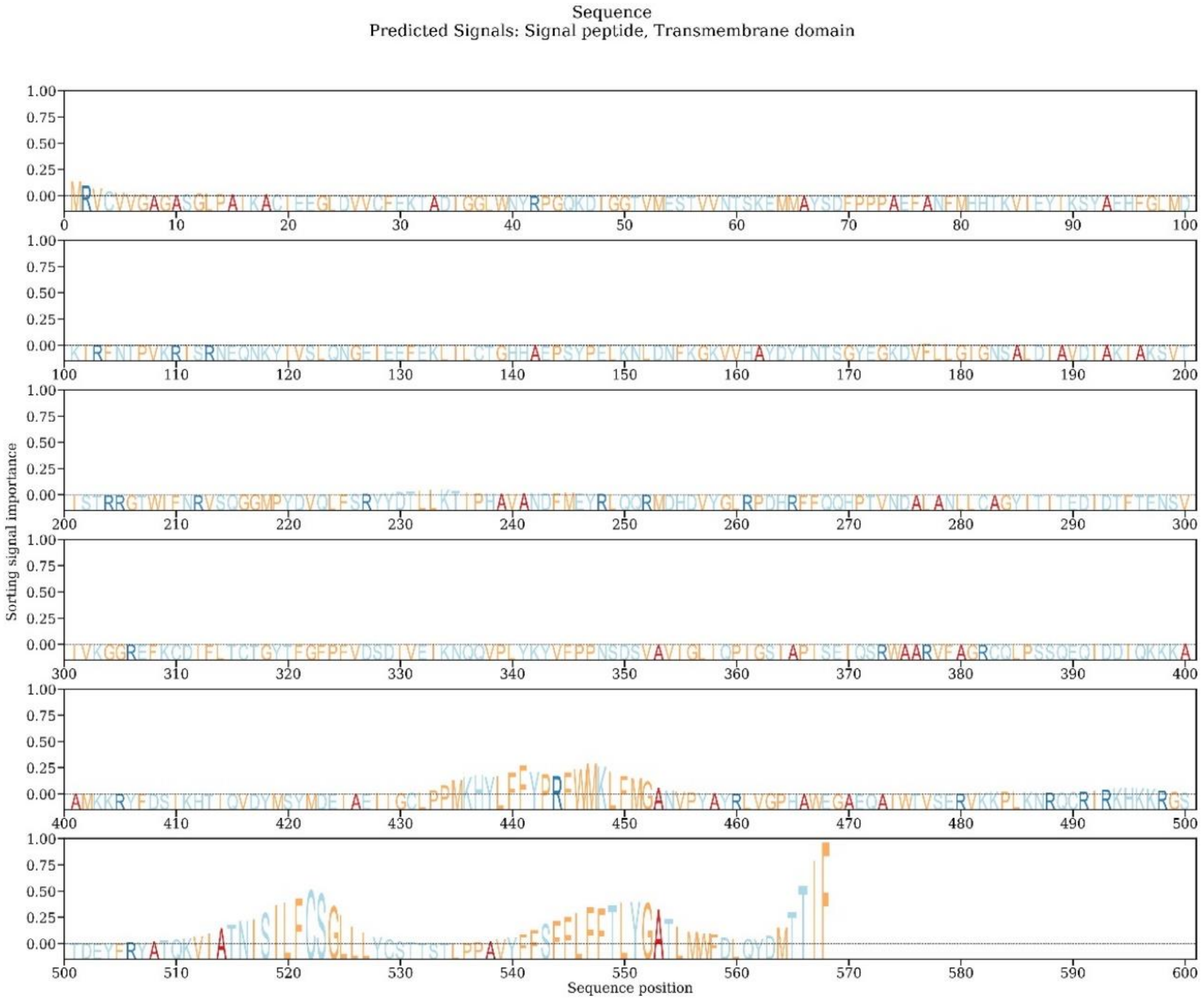


**Supplementary Figure 2. Subcellular localization prediction based on protein sequence of *C. elegans* FMO-4.** Based on protein sequence, *C. elegans* FMO-4 is predicted to be located in the endoplasmic reticulum and the golgi apparatus. Analysis was done using Deeploc-2.0.

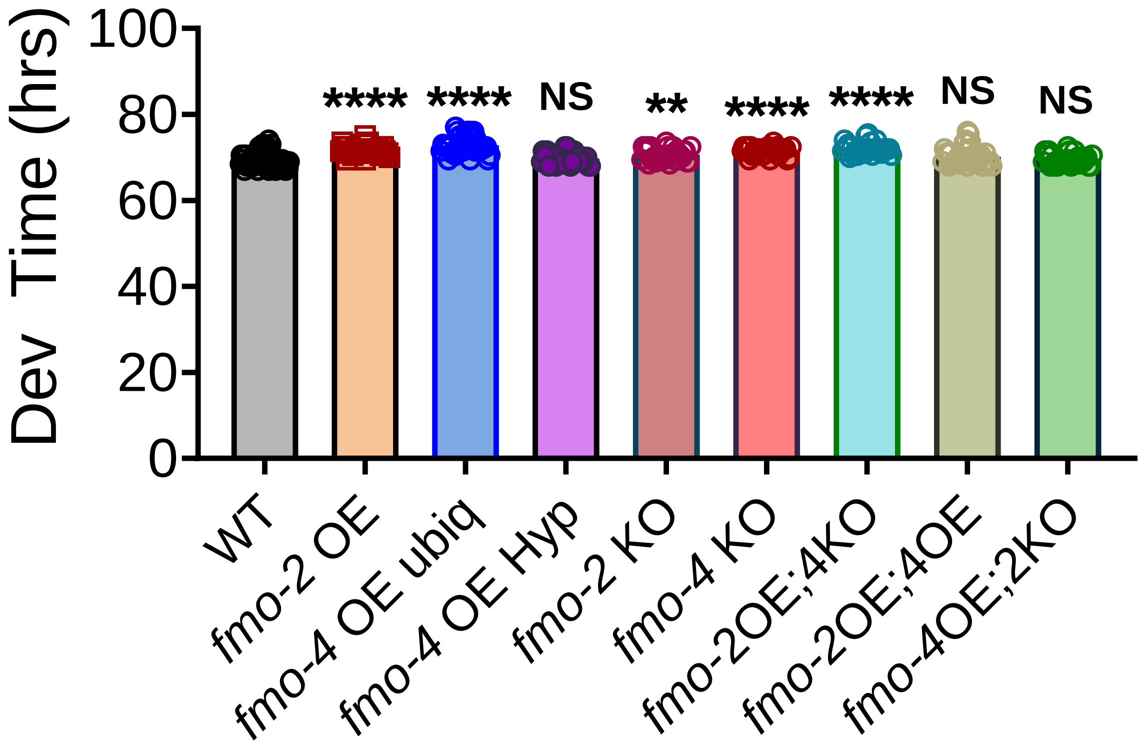


**Supplementary Figure 3. Development time of experimental worm strains compared to wild-type.** Development (Dev) time in hours (hrs) of wild-type (WT), *fmo-2* overexpressing (OE), ubiquitous *fmo-4* OE (*fmo-4* OE ubiq), hypodermal-specific *fmo-4* OE (*fmo-4* OE Hyp), *fmo-2* knockout (KO), *fmo-4* KO, *fmo-2* OE;*fmo-4* KO (*fmo-2*OE;*4*KO), *fmo-2* OE;*fmo-4* OE (*fmo-2*OE;*4*OE), and *fmo-4* OE;*fmo-2* KO (*fmo-4*OE;*2*KO) worms. (n = ~10 worms per condition, three replicate experiments). * denotes significant change in development time compared to WT. $p < 0.05$ using unpaired two-tailed t test. NS = not significant.

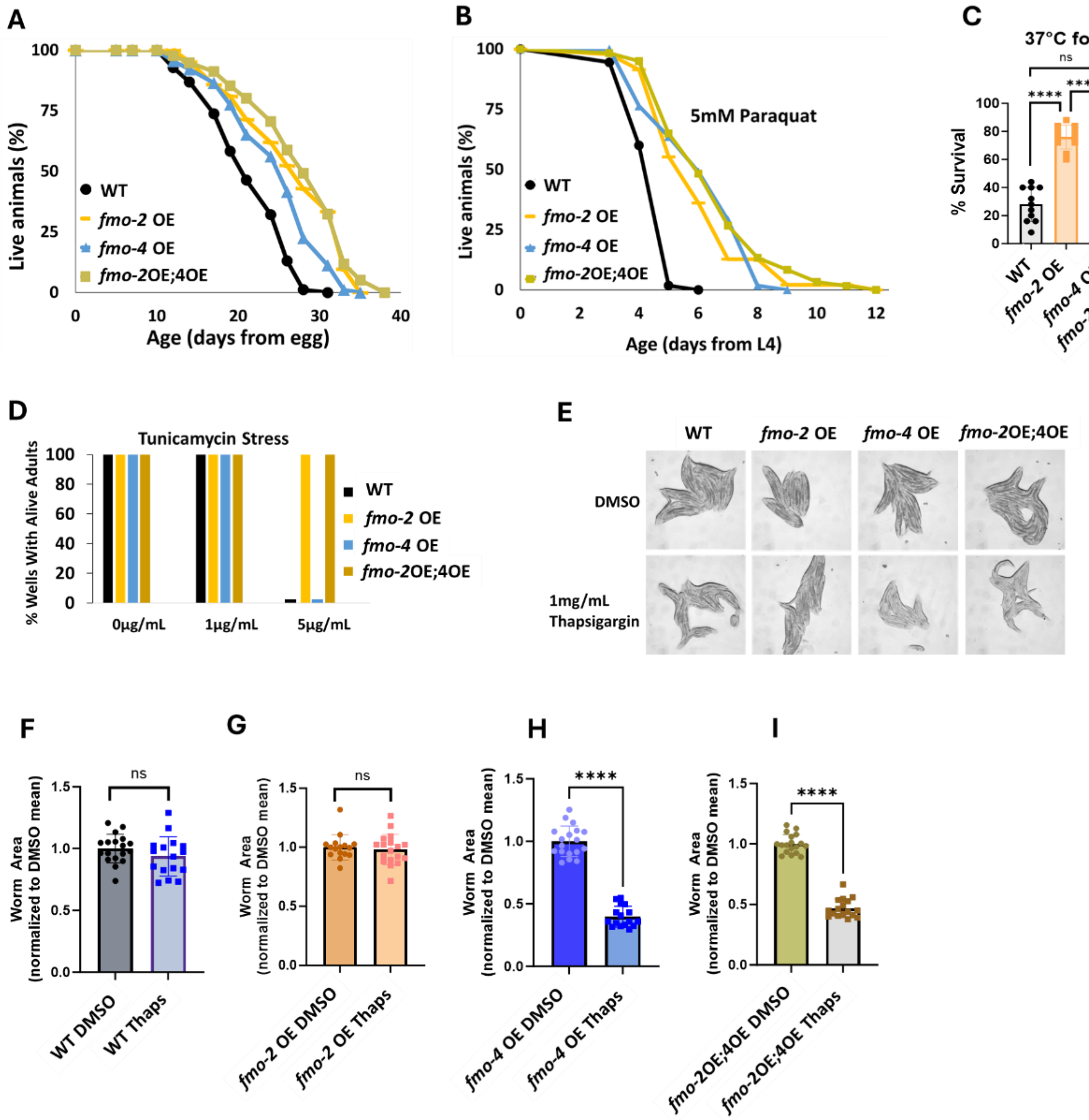


**Supplementary Figure 4. *fmo-4* acts in the same pathway as *fmo-2.*** (A) Lifespan analysis of wild-type (WT), *fmo-2* overexpressing (*fmo-2* OE), *fmo-4* overexpressing (*fmo-4* OE), and *fmo-2* OE;*fmo-4* OE (*fmo*-2OE;4OE) worms starting from egg (n = ~120 worms per condition, three replicate experiments). Significance was determined at $p < 0.05$ using log-rank analysis. (B) Survival of worms exposed to 5mM paraquat starting from L4 stage (n = ~90 worms per condition, three replicate experiments). Significance was determined at $p < 0.05$ using log-rank analysis. (C) Survival of worms exposed to 37°C heat for 3 hours (hrs) at L4 stage (n = 100 worms per condition, three replicate experiments). (D) Survival of worms exposed to 0, 1, and 5ug/mL tunicamycin starting from egg until day 1 of adulthood (n = ~60 eggs per condition, three replicate experiments). (E) Brightfield images of WT, *fmo-2* OE, *fmo-4* OE, and *fmo*-2OE;4OE worms exposed to DMSO or 1mg/mL thapsigargin (n = ~20 worms per condition, three replicate experiments). Quantification of (E) in (F-I). (J) For heat stress, tunicamycin stress, and thapsigargin stress, * denotes significant change at $p < 0.05$ using unpaired two-tailed t test or one-way ANOVA. N.S. = not significant.

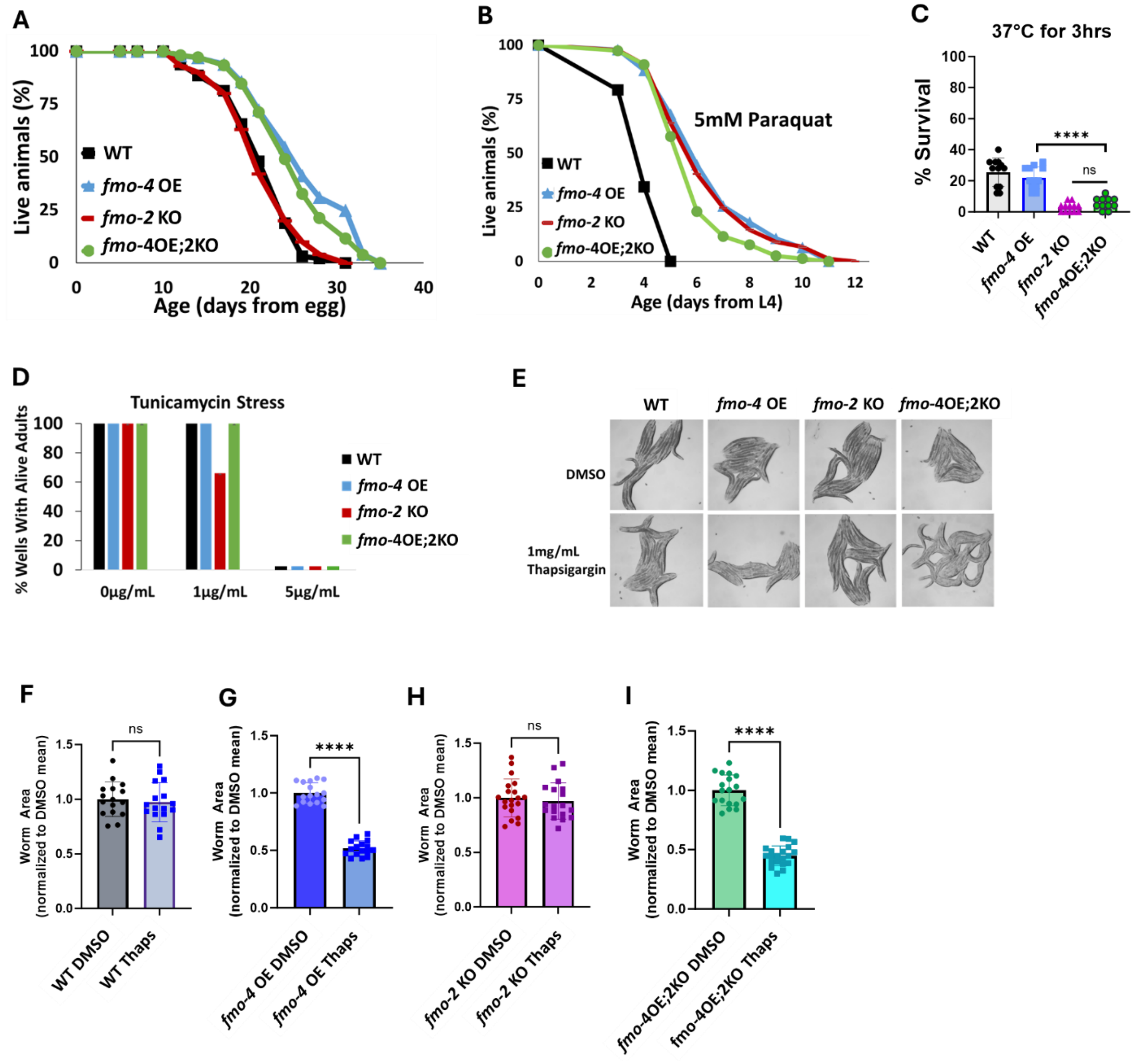


**Supplementary Figure 5. Knocking out *fmo-2* does not affect *fmo-4* overexpression.** (A) Lifespan analysis of wild-type (WT), *fmo*-4 overexpressing (*fmo-4* OE), *fmo-2* knockout (*fmo-2* KO), and *fmo-4* OE;*fmo-2* KO (*fmo-4*OE;*2*KO) worms starting from egg (n = ~120 worms per condition, three replicate experiments). Significance was determined at $p < 0.05$ using log-rank analysis. (B) Survival of worms exposed to 5mM paraquat starting at L4 stage (n = ~90 worms per condition, three replicate experiments). Significance was determined at $p < 0.05$ using log-rank analysis. (C) Survival of worms exposed to 37°C heat for 3 hours (hrs) from L4 stage (n = 100 worms per condition, three replicate experiments). (D) Survival of worms exposed to 0, 1, and 5ug/mL tunicamycin starting from egg until day 1 of adulthood (n = ~60 eggs per condition, three replicate experiments). (E) Brightfield images of WT, *fmo-4* OE, *fmo-2* KO, and *fmo-4*OE;*2*KO worms exposed to DMSO or 1mg/mL thapsigargin (n = ~20 worms per condition, three replicate experiments). Quantification of (E) in (F-I). For heat stress, tunicamycin stress, and thapsigargin stress, * denotes significant change at $p < 0.05$ using unpaired two-tailed t test or one-way ANOVA. N.S. = not significant.

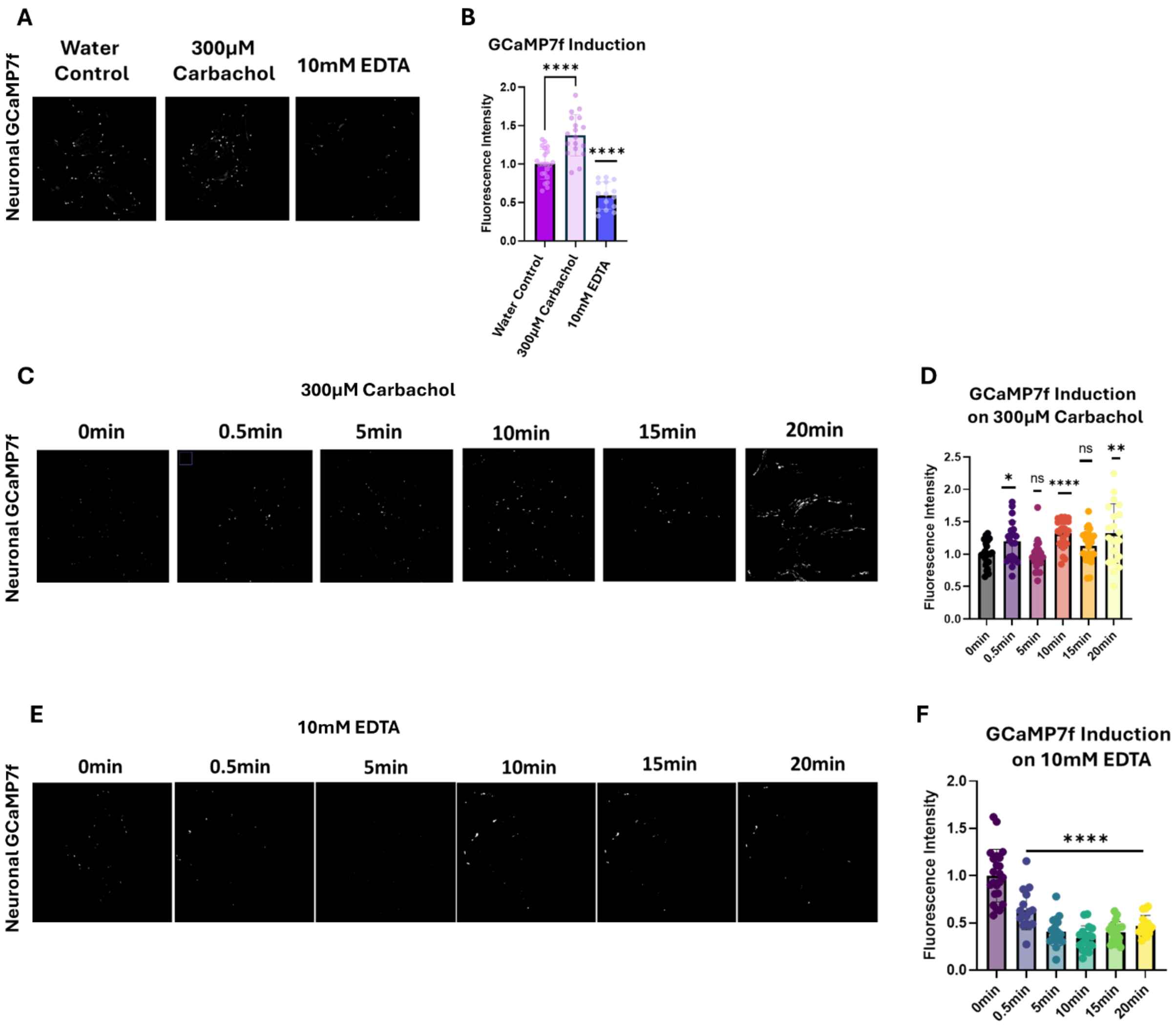


**Supplementary Figure 6. Carbachol and EDTA alter GCaMP7f fluorescence intesity.** (A) Fluorescence images of neuronal GCaMP7f calcium indicator worms on water control, 300µM carbachol, or 10mM EDTA. Fluorescence intensity is quantified in (B) (n = ~20 worms per condition, three replicate experiments). (C) Fluorescence images of the neuronal GCaMP7f calcium indicator worms on 300µM carbachol assessed at multiple time points including 0min (minutes), 0.5min, 5min, 10min, 15min, and 20min. Fluorescence intensity is quantified in (D) (n = ~20 worms per condition, three replicate experiments). (E) Fluorescence images of the neuronal GCaMP7f calcium indicator worms on 10mM EDTA assessed at multiple time points including 0 minutes (min), 0.5min, 5min, 10min, and 20min. Fluorescence intensity is quantified in (F) (n = ~20 worms per condition, three replicate experiments). * denotes significant change in fluorescence compared to control. $p < 0.05$ using unpaired two-tailed t test. N.S. = not significant.

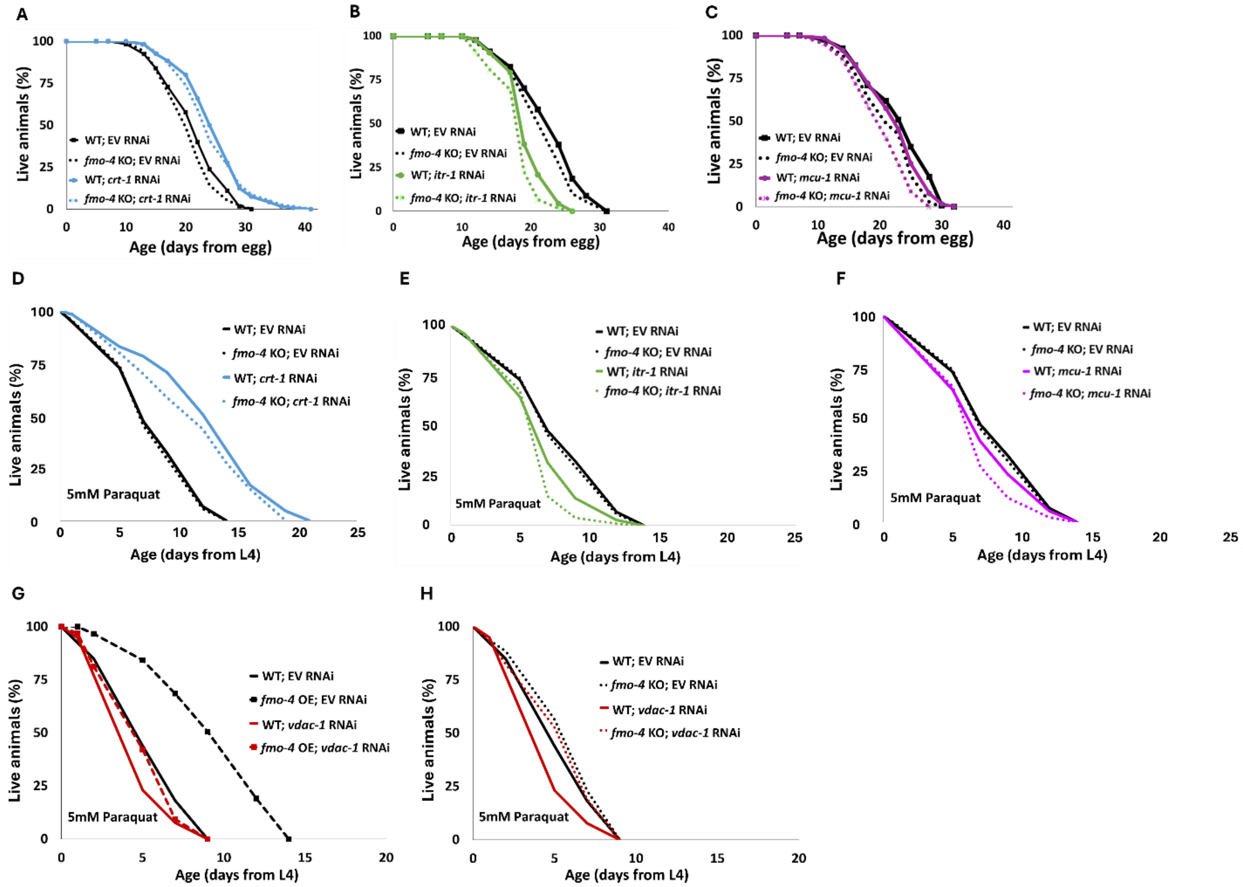


**Supplementary Figure 7. Lifespan and paraquat survival assessments of *fmo-4* KO worms on *crt-1, itr-1,* and *mcu-1* RNAi.** (A) Lifespan assessment of wild-type (WT) and *fmo-4* knockout (*fmo-4* KO) worms on empty vector (EV) and *crt-1* RNAi. (B) Lifespan assessment of WT and *fmo-4* KO worms on EV and *itr-1* RNAi. (C) Lifespan assessment of WT and *fmo-4* KO worms on EV and *mcu-1* RNAi. For all lifespan assays, n = ~120 worms per condition, three replicate experiments performed. (D) Survival of WT and *fmo-4* KO worms on EV and *crt-1* RNAi exposed to 5mM paraquat at L4 stage. (E) Survival of WT and *fmo-4* KO worms on EV and *itr-1* RNAi exposed to 5mM paraquat at L4 stage. (F) Survival of WT and *fmo-4* KO worms on EV and *mcu-1* RNAi exposed to 5mM paraquat at L4 stage. (G) Survival of WT and *fmo-4* overexpressing (OE) worms on EV and *vdac-1* RNAi exposed to 5mM paraquat at L4 stage. (H) Survival of WT and *fmo-4* KO worms on EV and *vdac-1* RNAi exposed to 5mM paraquat at L4 stage. For all paraquat survival assays, n = ~90 worms per condition, three replicate experiments performed. Significance was determined at $p < 0.05$ using log-rank analysis and significant interactions between the condition of interest and genotype was determined at $p < 0.01$ using Cox regression analysis.

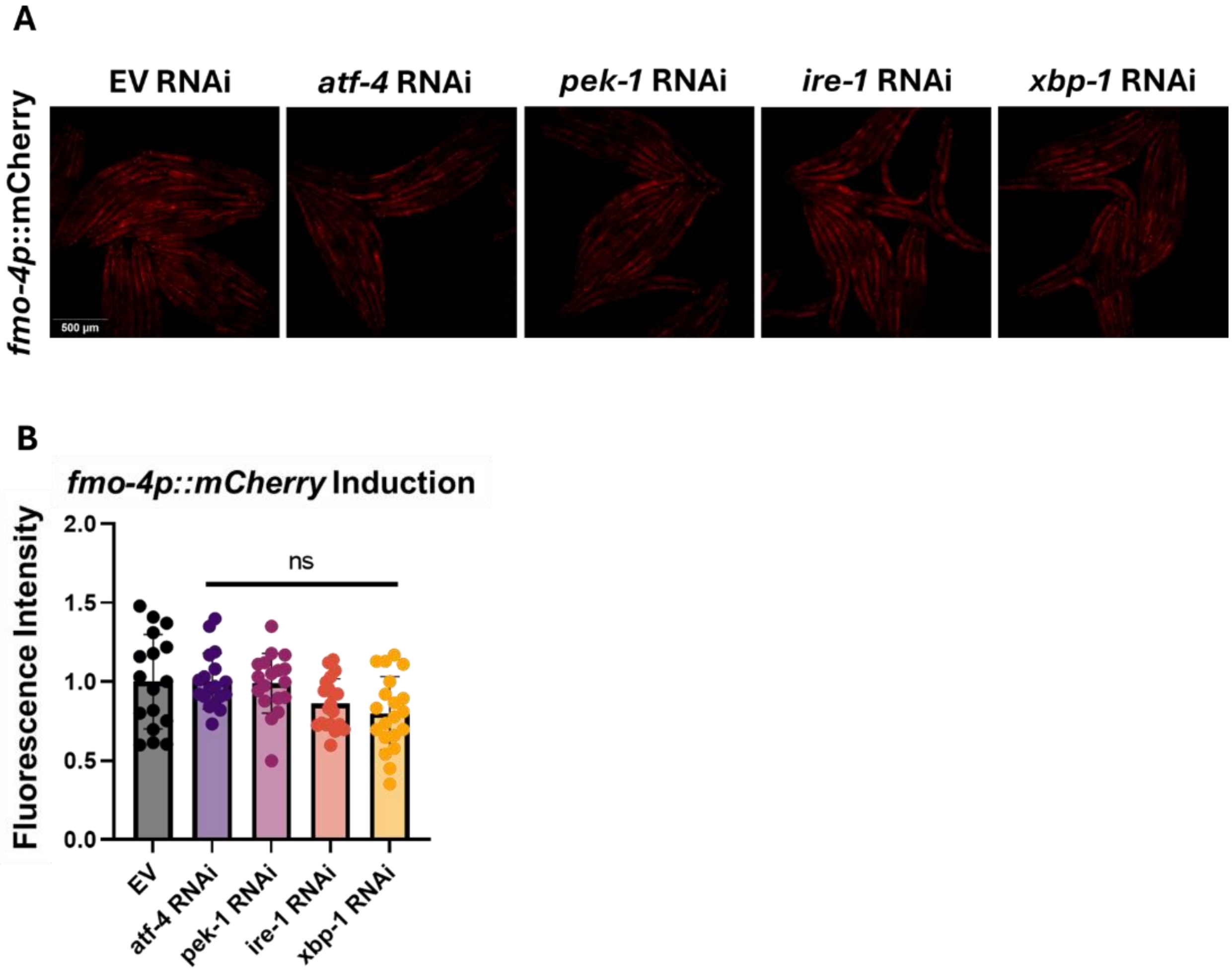


**Supplementary Figure 8. *fmo-4* gene expression is not induced by the other branches of UPR$^{ER}$.** (A) Fluorescence images of the *fmo-4p::mCherry* reporter worms on empty vector (EV) RNAi, *atf-4* RNAi, *pek-1* RNAi, *ire-1* RNAi, and *xbp-1* RNAi, quantified in (B) (n = ~20 worms per condition, three replicate experiments). * denotes significant change in fluorescence compared to EV RNAi control. $p < 0.05$ using unpaired two-tailed t test. N.S. = not significant.

**Supplementary Table 1. qPCR validation of worm strains.**

| Worm Strain | qPCR Primer | Fold Change | Replicate # |
|---|---|---|---|
| *fmo-2* OE;4KO | *fmo-2* | 51 | 1 |
| *fmo-2* OE;4KO | *fmo-2* | 50.7 | 2 |
| *fmo-2* OE;4KO | *fmo-2* | 45.7 | 3 |
| *fmo-2* OE;4KO | *fmo-4* | 0.00007 | 1 |
| *fmo-2* OE;4KO | *fmo-4* | 0.00043 | 2 |
| *fmo-2* OE;4KO | *fmo-4* | 0.00005 | 3 |
| *fmo-2* OE;4OE | *fmo-2* | 82.6 | 1 |
| *fmo-2* OE;4OE | *fmo-2* | 152.7 | 2 |
| *fmo-2* OE;4OE | *fmo-2* | 63.6 | 3 |
| *fmo-2* OE;4OE | *fmo-4* | 39 | 1 |
| *fmo-2* OE;4OE | *fmo-4* | 42.1 | 2 |
| *fmo-2* OE;4OE | *fmo-4* | 29.3 | 3 |
| *fmo-* 4OE;2KO | *fmo-2* | 0.009 | 1 |
| *fmo-* 4OE;2KO | *fmo-2* | 0.03 | 2 |
| *fmo-* 4OE;2KO | *fmo-2* | 0.002 | 3 |
| *fmo-* 4OE;2KO | *fmo-4* | 39 | 1 |
| *fmo-* 4OE;2KO | *fmo-4* | 76 | 2 |
| *fmo-* 4OE;2KO | *fmo-4* | 45.5 | 3 |
| *fmo* -4OE ubiq | *fmo-4* | 139.7 | 1 |
| *fmo* -4OE ubiq | *fmo-4* | 164.4 | 2 |
| *fmo* -4OE ubiq | *fmo-4* | 94.41 | 3 |
| *fmo* -4OE$^{Hyp}$ | *fmo-4* | 44.51 | 1 |
| *fmo* -4OE$^{Hyp}$ | *fmo-4* | 46.6 | 2 |
| *fmo* -4OE$^{Hyp}$ | *fmo-4* | 44.4 | 3 |
| *fmo* -4 KO | *fmo-4* | 0.00003 | 1 |
| *fmo* -4 KO | *fmo-4* | 0.000003 | 2 |
| *fmo* -4 KO | *fmo-4* | 0.00004 | 3 |

**Supplementary Table 2. DAVID analysis shows that calcium ion binding transcripts are regulated when *fmo-4* is overexpressed in *C. elegans.***

| Gene ID | Gene Name | Species |
|---|---|---|
| WBGene00018304 | Agrin (*agr-1* ) | Caenorhabditis elegans |
| WBGene00001475 | Cadherin EGF LAG seven-pass G-type receptor fmi-1 (*fmi-1* ) | Caenorhabditis elegans |
| WBGene00022103 | Cadherin domain-containing protein (*cdh-12* ) | Caenorhabditis elegans |
| WBGene00000401 | Cadherin domain-containing protein (*cdh-9* ) | Caenorhabditis elegans |
| WBGene00000396 | Cadherin-4 (*cdh-4* ) | Caenorhabditis elegans |
| WBGene00017028 | Dendrite extension defective protein 1 (*dex-1* ) | Caenorhabditis elegans |
| WBGene00019184 | EF-hand domain-containing protein (H10E21.4) | Caenorhabditis elegans |
| WBGene00019285 | EF-hand domain-containing protein (*cbn-1* ) | Caenorhabditis elegans |
| WBGene00008779 | EGF-like domain-containing protein (F14B4.1) | Caenorhabditis elegans |
| WBGene00022816 | EGF-like domain-containing protein (*fbn-1* ) | Caenorhabditis elegans |
| WBGene00013416 | Multiple epidermal growth factor-like domains protein 6 (Y64G10A.7) | Caenorhabditis elegans |
| WBGene00003772 | Neurexin like receptor 1 (*nlr-1* ) | Caenorhabditis elegans |
| WBGene00007170 | Nidogen (B0393.5) | Caenorhabditis elegans |
| WBGene00000792 | uncharacterized protein (*crb-1* ) | Caenorhabditis elegans |

**Supplementary Table 3. List of RNAi used.**

| RNAi | Description | E. coli strain | RNAi Library |
|---|---|---|---|
| EV | Empty Vector RNAi control | HT115 | Ahringer |
| *crt-1* | calreticulin knock down | HT115 | Ahringer |
| *itr-1* | inositol triphosphate receptor knock down | HT115 | Ahringer |
| *mcu-1* | mitochondrial calcium uniporter knock down | HT115 | Ahringer |
| *atf-6* | activating transcription factor-6 knock down | HT115 | Ahringer |
| *atf-4* | activating transcription factor-4 knock down | HT115 | Ahringer |
| *pek-1* | PERK kinase knock down | HT115 | Ahringer |
| *ire-1* | IRE1 kinase knock down | HT115 | Ahringer |
| *xbp-1* | X-box binding protein knock down | HT115 | Ahringer |

**Supplementary Table 4. List of worm strains used in this paper.**

| Worm Strain Description | Source | Identifier |
|---|---|---|
| Wild-Type | CGC | N2 or WT |
| *fmo-4(ok294)* | CGC | *fmo-4* KO |
| *fmo-2(ok2147)* | CGC | *fmo-2* KO |
| *eft-3p::fmo-4::* SL2-GFP-let-858 3'UTR | Suny BioScience | *fmo-4* OE ubiquitous |
| *dpy-7p::fmo-4* ::SL2-GFP-let-858 3'UTR | Suny BioScience | *fmo* -4 OE hypodermal |
| KAE9; *fmo-4(ok294)* | In House | *fmo* -2OE;4KO |
| KAE9; LEI168 | In House | *fmo* -2OE;4OE |
| *eft-3p::fmo-4* ::SL2-GFP-let-858 3'UTR;*fmo-2(ok2147)* | In House | *fmo* -4OE;2KO |
| *eft-3p::fmo-2* + *h2b ::gfp* + *Cbr-unc-119(+)* | Injection In House | *fmo* -2 OE |
| *fmo-4p::mCherry* | Suny BioScience | *fmo-4p::mCherry* |

**Supplementary Table 5. List of genotyping primers used to validate worm strains.**

| Genotyping Primer | Sequence |
|---|---|
| *fmo-4_ok294_* F (*fmo-4* KO) | GCATATGGTACATTTGCCCC |
| *fmo-4_ok294_* R (*fmo-4* KO) | AAATCAACTCAAACCGCACC |
| *fmo-2_ok2147_* F (*fmo-2* KO) | TGTTCAATGATCGTACCCGA |
| *fmo-2_ok2147_* R (*fmo-2* KO) | GTTCGAAATTGGGAAATCCA |
| *ttTi4348_ChrI_HR_* F (*fmo-2* OE) | AATTCCTGAGCTGCATGGCT |
| *ttTi4348_ChrI_HR_* R (*fmo-2* OE) | GTCGACCGCTAGTGTAGCTT |
| *ttTi4348_ChrI_internal_* R (*fmo-2* OE) | TGAGCCAATTCATCCCGGTT |
| *fmo-4* OE ubiquitous | visualized green fluorescence |
| *fmo-4* OE hypodermal | visualized green fluorescence |

**Supplementary Table 6. List of qPCR primers used to validate worm strains.**

| qPCR Primer | Sequence |
|---|---|
| *cdc-42* _qPCR_F | CTGCTGGACAGGAAGATTACG |
| *cdc-42* _qPCR_R | CTCGGACATTCTCCAATGAAG |
| Y45F10D4_qPCR_F | GTCGCTTCAAATCAGTTCAGC |
| Y45F10D4_qPCR_R | GTTCTTGTCAAGTGATCCGACA |
| *fmo-2* _qPCR_F | ACGAAACGAATGAGTCGTCAGT |
| *fmo-2* _qPCR_R | AGAGCAGACAAGAACGCCAT |
| *fmo-4* _qPCR_F | AGGGAGGCAGAGAATTCAAGT |
| *fmo-4* _qPCR_R | GGGAAATCCGAGTAGGCC |
